# Physiological responses of two accessions of tomato to three distance regimes in juvenile oil palm-tomato intercrop in rainforest zone of nigeria

**DOI:** 10.1101/2020.05.31.126771

**Authors:** Ayodele Samuel Oluwatobi

## Abstract

This study was conducted during the rainy season of 2017 within the alleys of juvenile oil palms that were 2-year-old at the oil palm plantation located in Ala, Akure-North Local Government, Ondo State. Tomato accessions (NGB 01665 and NG/AA/SEP/09/053) were intercropped at 1, 2 and 3 m from the juvenile oil palm within the alley in a randomized complete block design. The results showed that tomato (NGB 01665 and NG/AA/SEP/09/053) planted at 3 m from the juvenile oil palm performed better than those at other planting distances in terms of growth and yield attributes with number of fruit; fruit weight and yield per hectare of 39.50, 2265.8 g and 3.74 ton/ha respectively. However, the control (sole) recorded the best yield but was not significantly different from those planted at 3 m from the juvenile oil palm. Varietal advantage was recorded by tomato (NGB 01665) with higher number of fruits, fruit weight and yield per hectare (26.94, 1834 g and 3.158 ton/ha) respectively. Intercropping advantage was not recorded for any of the intercropping distance regimes during the trial (when the juvenile oil palms were 2-year-old), with LER values less than unity.

## INTRODUCTION

The oil palm (*Elaeis guineensis* Jacq.) is one of the important economic crops in the tropics (Anyanwu *et al.*, 1982; Ibitoye *et al.*, 2011). Lack of modern farm mechanization, over dependency on smallholder/traditional processors, land tenure problem, inadequate infrastructure, poor funding, campaign by environmentalist for environmental protection etc. are some of the identified problems that have limited the cultivation and production of oil palm in Nigeria (Sridhar and Adeoluwa, 2009). Soyebo *et al*. (2005) reported that land is the major factor limiting oil palm cultivation. Their report recognized that majority of the farmers in Nigeria (81%) are confronted with land problem, 34.2% with fund problem while 53.2% complained of inadequate information and cultivation knowledge about oil palm. The standard 9 × 9 m triangular spacing use for oil palm provides wide spaces between the young palms (Okigbo, 1979). This leads to considerable waste of solar radiation and weed problem from transplanting to canopy closure which takes between three to five years (Chee *et al*., 2000). The only way to increase agricultural production in the small or marginal units of farming is to increase the productivity per unit time and area (Chatterjee *et al.*, 1993). Growing a number of other food crops in association with juvenile oil palm trees is a widespread practice in most oil palm growing areas in Nigeria. Intercropping vegetables with different architecture and nutritional value such as beet and okra, pepper and onion with juvenile oil palm has been practised in tropical Asia (Okigbo, 1979). Nuertey *et al*. (2000) identified a number of crops that the farmers intercrop with oil palm and the basis of their selection. Farmers may seem justified then by growing food and /or cash crops at different intercropping distances from the juvenile oil palm trees until canopy closure.

Therefore, the aim of this study is to investigate the performance of two accessions of tomato (NGB 01665 and NG/AA/SEP/09/053) when intercropped with juvenile oil palm at varying distances from the juvenile oil palm. The objectives of this study are to evaluate the growth and yield responses of the two tomato accessions to three spacing regimes in juvenile oil palm-tomato intercrop.

## MATERIALS AND METHODS

### Description of study area

The field experiment was conducted within an established juvenile oil palm plantation located at Ala in Akure-North Local Government Area of Ondo State (coordinate of 7.093 ° N, 5.354 °E). It is located in the tropical rain forest region of Nigeria; has predominant climatic characteristics of warm and humid with little seasonal variation. Annual rainfall varies from 1150 to 2550 mm. Temperature is moderately high year-round and range between 22 and 34 °C with daily average of about 30 °C (Ogunrayi *et al.*, 2016).

### Collection of tomato seeds

Two accessions of tomato seeds were obtained from National Centre for Genetic Resources and Biotechnology (NACGRAB) research institute, Ibadan Oyo State. The tomato accessions are: NGB 01665 and NG/AA/SEP/09/053. The tomato seeds were raised in a nursery for 4 weeks before transplanting into the alley of juvenile oil palm of 2 years old.

### Field experiment

The oil palm plantation has a spacing of 6 × 6m; the tomato seeds were transplanted at 1, 2 and 3 m from the juvenile oil palm. The experiment was laid out in a randomized complete block design.

Partial Land Equivalent Ratio (LER) of the tomato per treatment was evaluated by dividing the intercrop yield of the vegetable crop by the sole yield (control). 

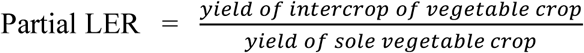

### Data collection

Data of growth parameters collected at 3 and 4 weeks after intercropping (WAI) include number of leaves, plant height, stem girth, leaf area, chlorophyll index, total chlorophyll content and dry matter. Number of leaves was obtained by physical counting of leaves; plant height was measured using a measuring tape; stem girth was measured using a vernier caliper; leaf area was measures using Easy leaf Area software for android devices (Easlon and Arnold, 2014); chlorophyll index was determined using handheld chlorophyll meter (atLEAF); total chlorophyll content was calculated by converting chlorophyll index values obtained from the handheld chlorophyll meter (atLEAF) at www.atleaf.com/conversion.

Reproductive parameters collected include number of fruits and yield per plant, weight of 100 fruits, fruit girth and length, and yield per hectare. Number of fruit per plant was determined by counting the number of fruits collected; weight of fruit per plant was determined by weighing the fruits collected from each tomato under the different spacing treatment; weight of 100 fruits was determined by weighing 100 fruits; fruit girth and length were determined with vernier caliper; and yield per hectare is determined by computing total weight of fruit per treatment in tonnage per hectare (Seran and Brintha, 2009).

Data analysis: Data collected from the study were analyzed using statistical package for social sciences (SPSS: version 16.0) by subjecting the data to analysis of variance (ANOVA). The mean values of the data were separated using Duncan Multiple Range Test (DMRT) at 5% probability level. Interaction analysis between the tomato accessions was carried out using SPSS. The results were presented in tables and charts. Graphs were plotted using Origin (version 7.0) software.

## RESULTS AND DISCUSSION

### Effect of intercropping on growth and yield components of tomato (NGB 01665)

In this present study, tomatoes of the control plot recorded the best performance in growth by recording highest values of leaf number, plant height and stem girth, however; were not significantly different from those planted at 3m from juvenile oil palm as shown in Figures 1-3.

**FIG. 1A-B:**
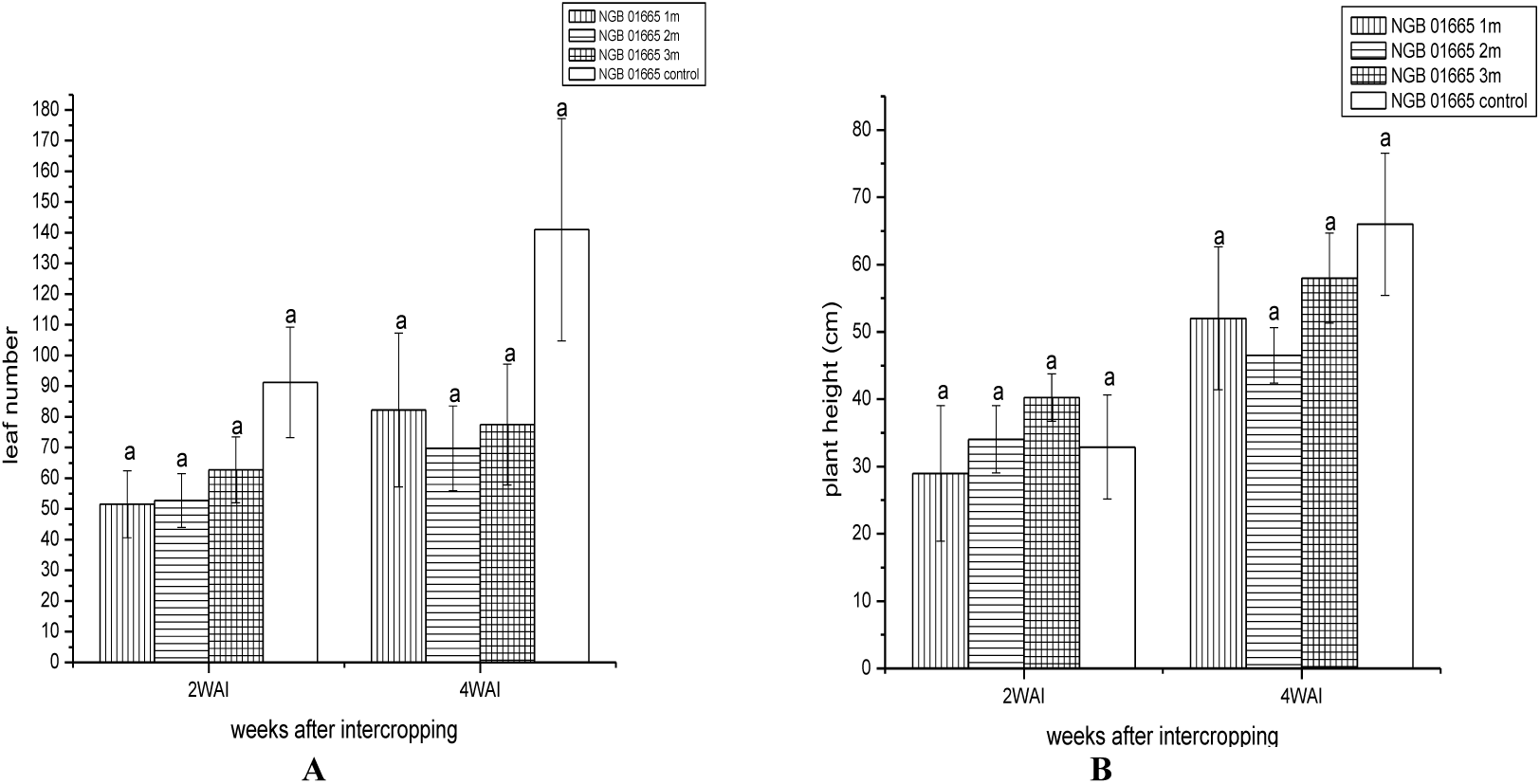
Effects of intercropping on the number of leaves of tomato (NGB 01665) at 2 and 4 WAI; effects of intercropping on the plant height of tomato (NGB 01665) at 2 and 4 WAI.

**FIG. 2A-B:**
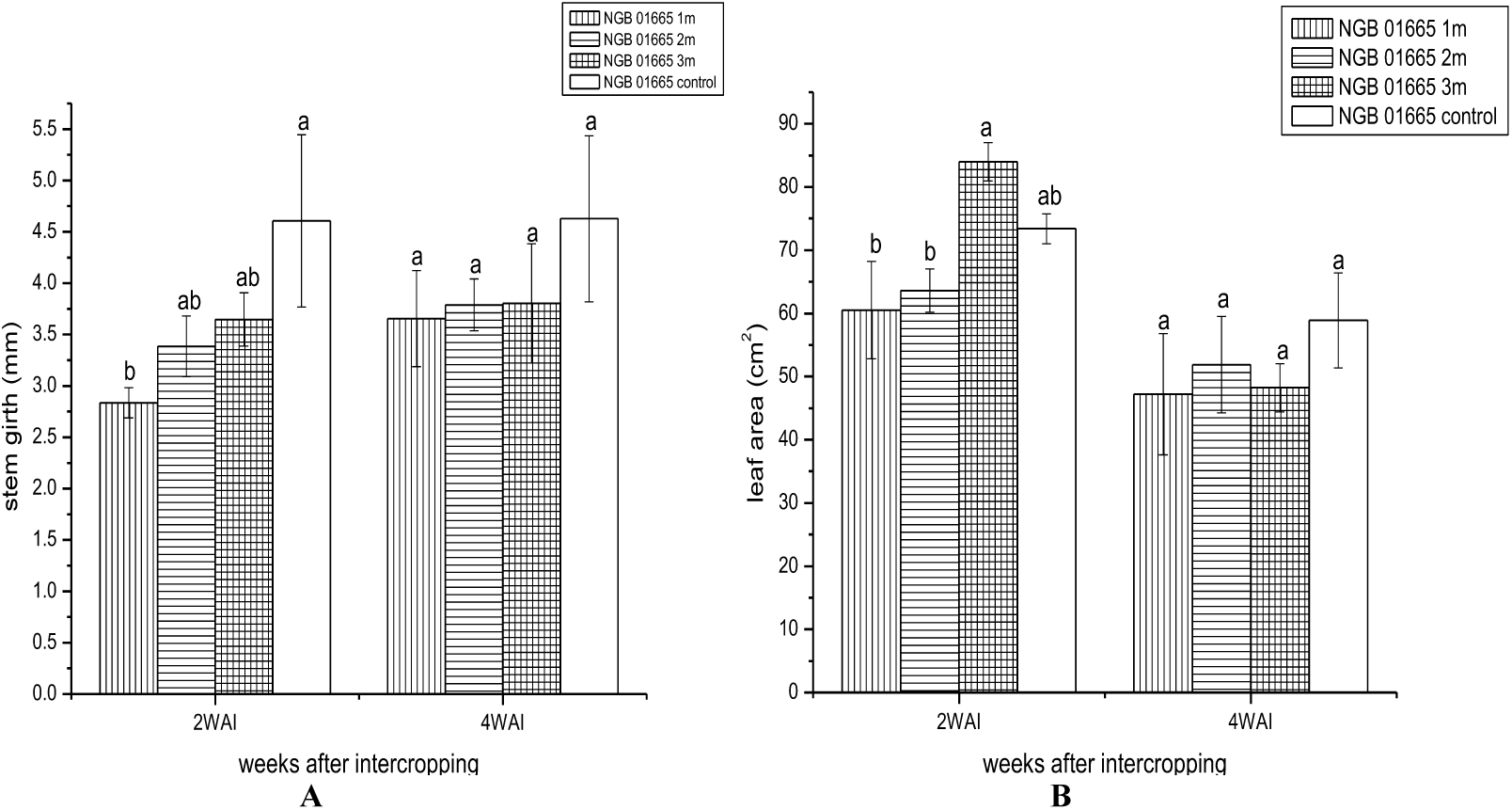
Effects of intercropping on the stem girth of tomato (NGB 01665) at 2 and 4 WAI; effects of intercropping on the leaf area of tomato (NGB 01665) at 2 and 4 WAI.

**FIG. 3A-B:**
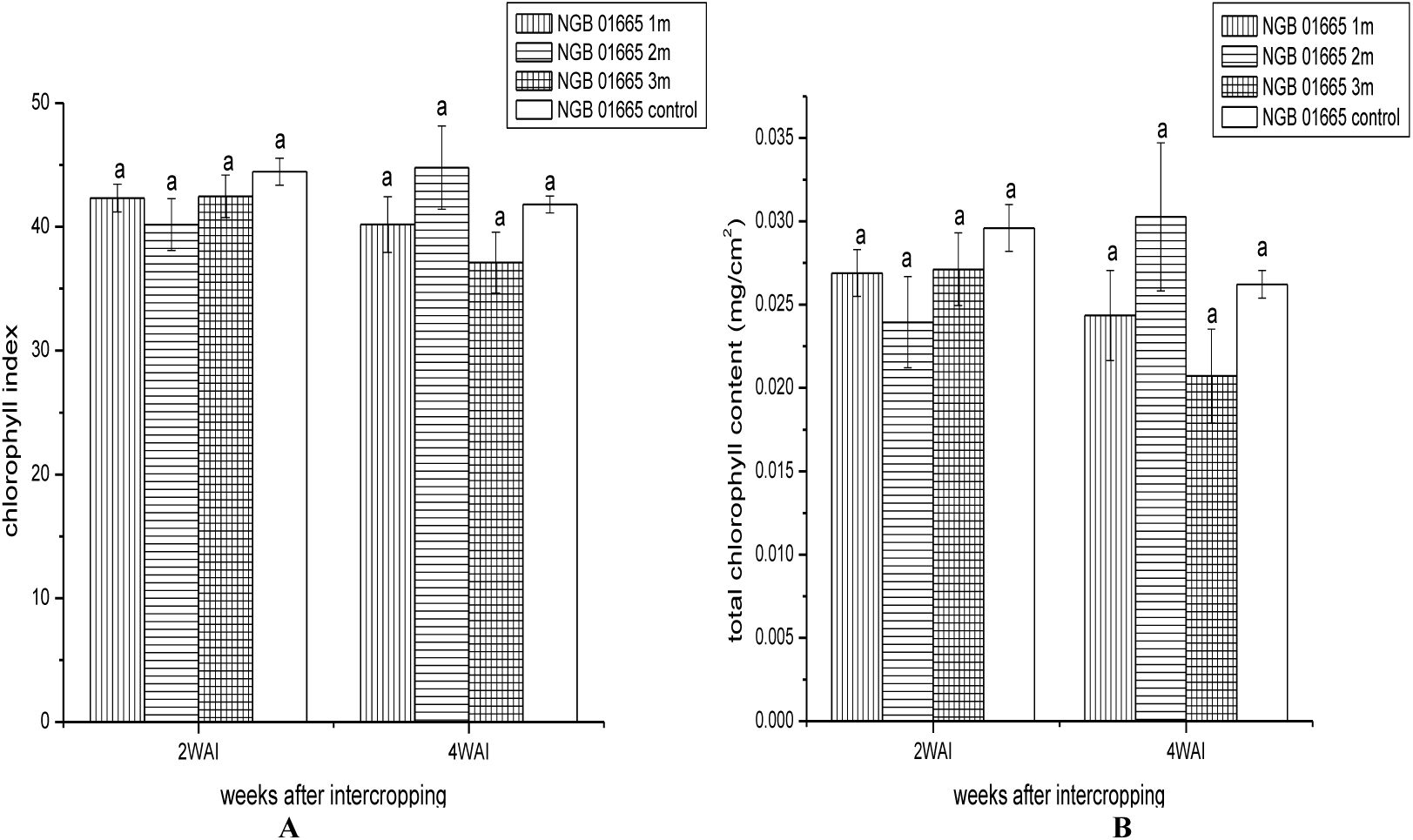
Effects of intercropping on the chlorophyll index of tomato (NGB 01665) at 2 and 4 WAI; effects of intercropping on total chlorophyll content of tomato (NGB 01665) at 2 and 4 WAI.

### Growth responses of tomato (NG/AA/SEP/09/053) to intercropping with juvenile oil palm

**FIG. 4A-B:**
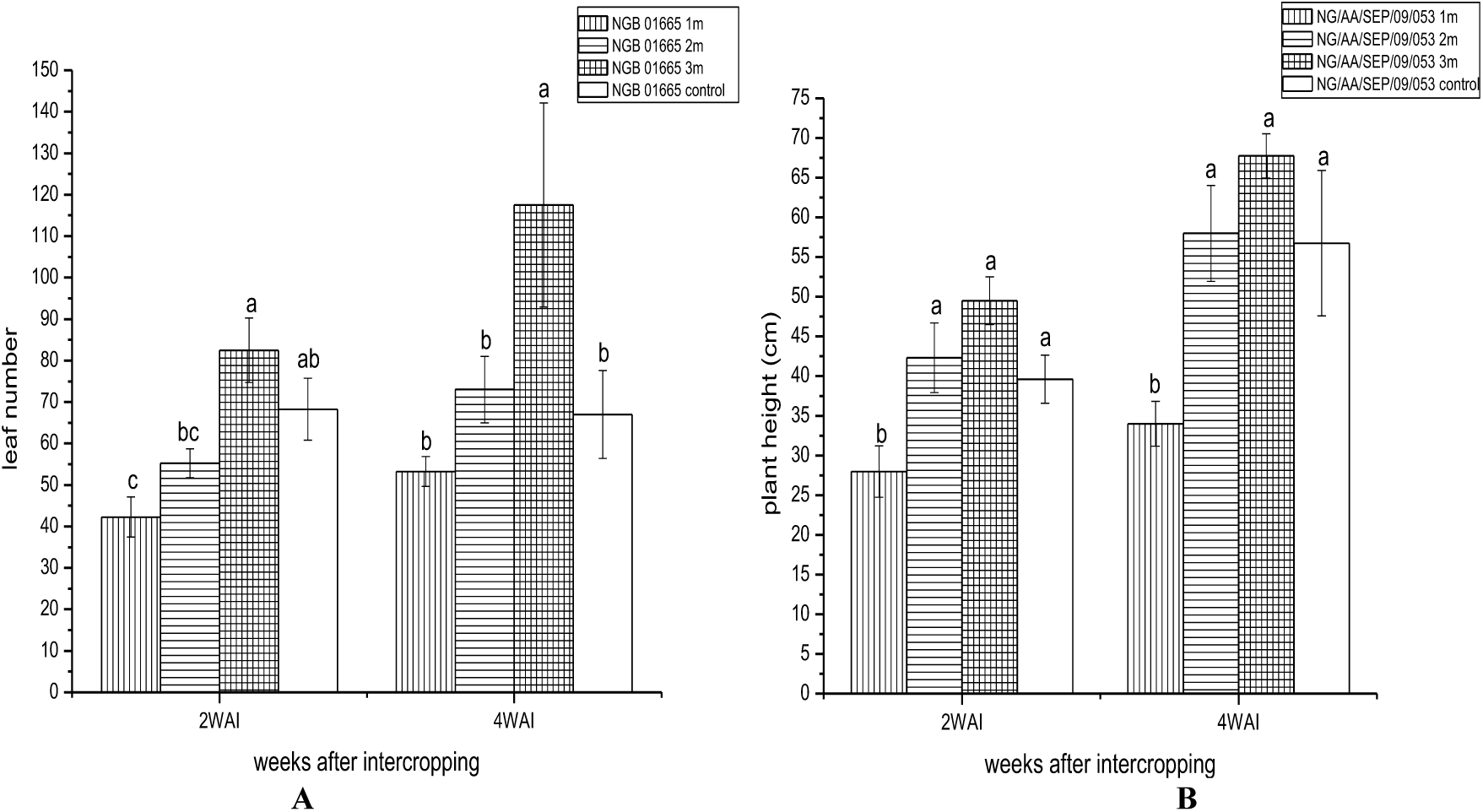
Effects of intercropping on number of leaves of tomato (NG/AA/SEP/09/053) at 2 and 4 WAI; effects of intercropping on plant height of tomato (NG/AA/SEP/09/053) at 2 and 4 WAI.

**FIG. 5A-B:**
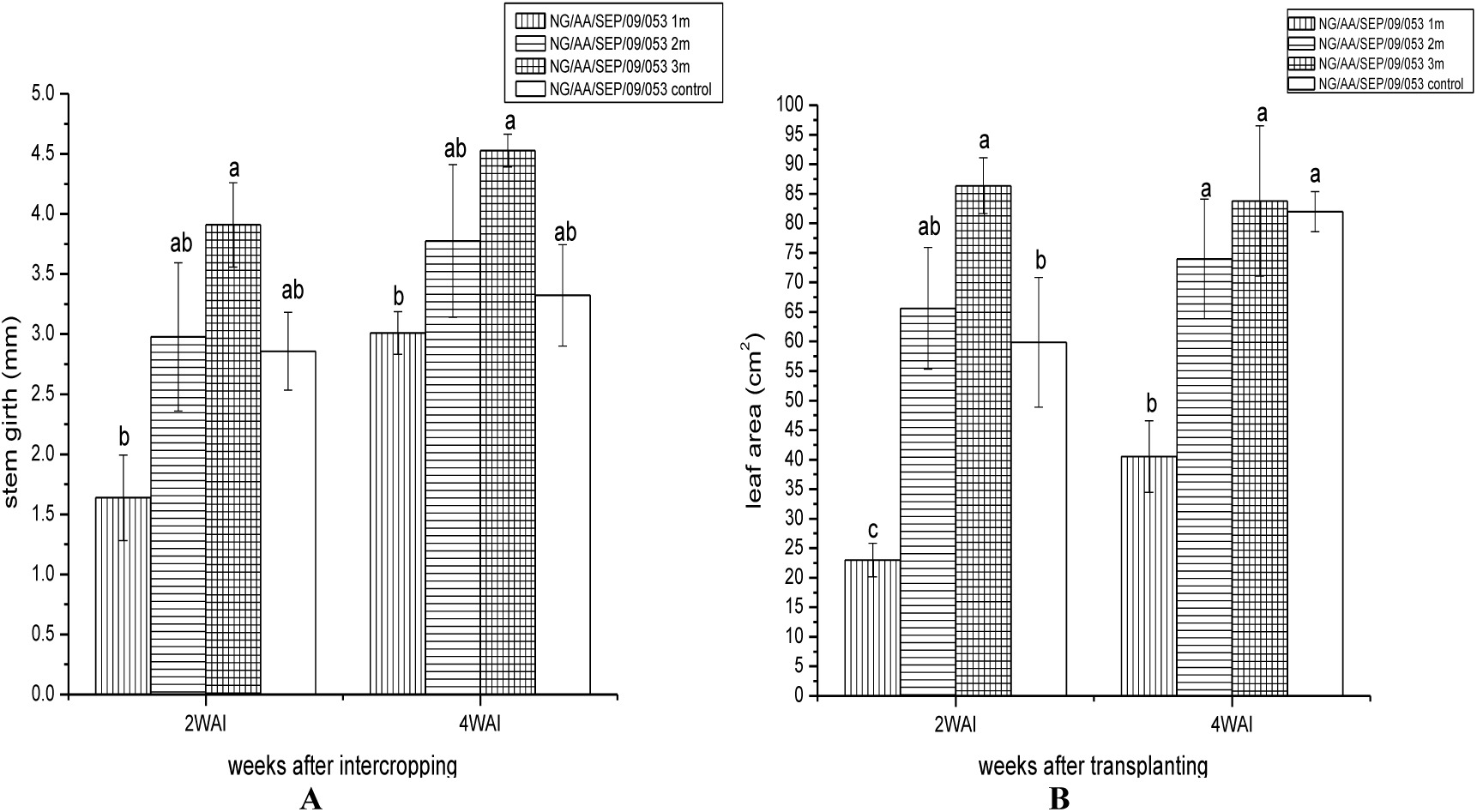
Effects of intercropping on stem girth of tomato (NG/AA/SEP/09/053) at 2 and 4 WAI; effects of intercropping on leaf area of tomato (NG/AA/SEP/09/053) at 2 and 4 WAI.

**FIG. 6A-B:**
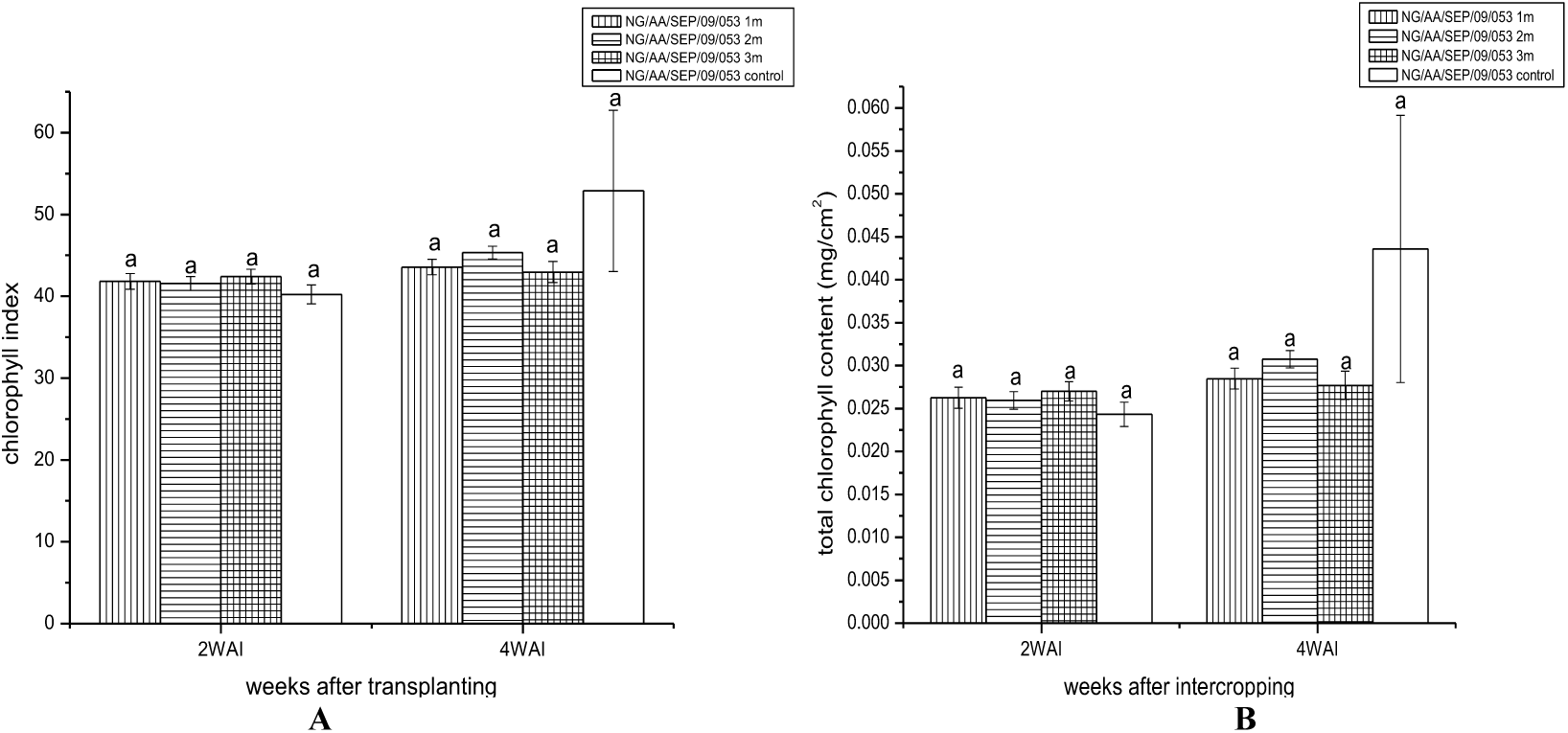
Effects of intercropping on index of tomato (NG/AA/SEP/09/053) at 2 and 4 WAI; effects of intercropping on total chlorophyll content of tomato (NG/AA/SEP/09/053) at 2 and 4 WAI.

These results indicate that tomato (NGB 01665) has high requirement for sunshine as the partial shade casted on those at 1 m and 2 m from juvenile oil palm could be responsible for the poor performances recorded at these planting distances. Njoroge and Kimemia (1995) worked on the economic benefits of intercropping young Arabica and Robusta coffee with food crops, they reported that intercropping tomatoes among other crops were found to be economically viable.

The results of the yield and yield components in this trial indicated that tomato (NGB 01665) planted at the control plot gave the best performance however it was not significantly different from those intercropped at 3 m from the juvenile oil palm (P<0.05) as shown in Table 1.

**TABLE 1:** Effects of intercropping on yield components of tomato (NGB 01665)-juvenile oil palm intercrop.

| Treatment | Number of fruits/plant | Fruit weight/plant (g) | Weight of 100 fruits (g) | Fruit girth (mm) | Fruit length (mm) | Yield (tonha<sup>-1</sup>) |
| --- | --- | --- | --- | --- | --- | --- |
| 1 m | 12.25<sup>b</sup> | 882.38<sup>c</sup> | 6431.60<sup>a</sup> | 40.23<sup>b</sup> | 39.83<sup>b</sup> | 2.68<sup>b</sup> |
| 2 m | 18.75<sup>b</sup> | 2033<sup>b</sup> | 7179.40<sup>b</sup> | 42.59<sup>b</sup> | 48.90<sup>a</sup> | 3.08<sup>ab</sup> |
| 3 m | 39.50<sup>a</sup> | 2265.8<sup>b</sup> | 8847.80<sup>ab</sup> | 52.23<sup>a</sup> | 54.73<sup>a</sup> | 3.74<sup>ab</sup> |
| Control | 42.50<sup>a</sup> | 3172.8<sup>a</sup> | 10625.00<sup>a</sup> | 52.97<sup>a</sup> | 54.15<sup>a</sup> | 4.35<sup>a</sup> |

**TABLE 2:**
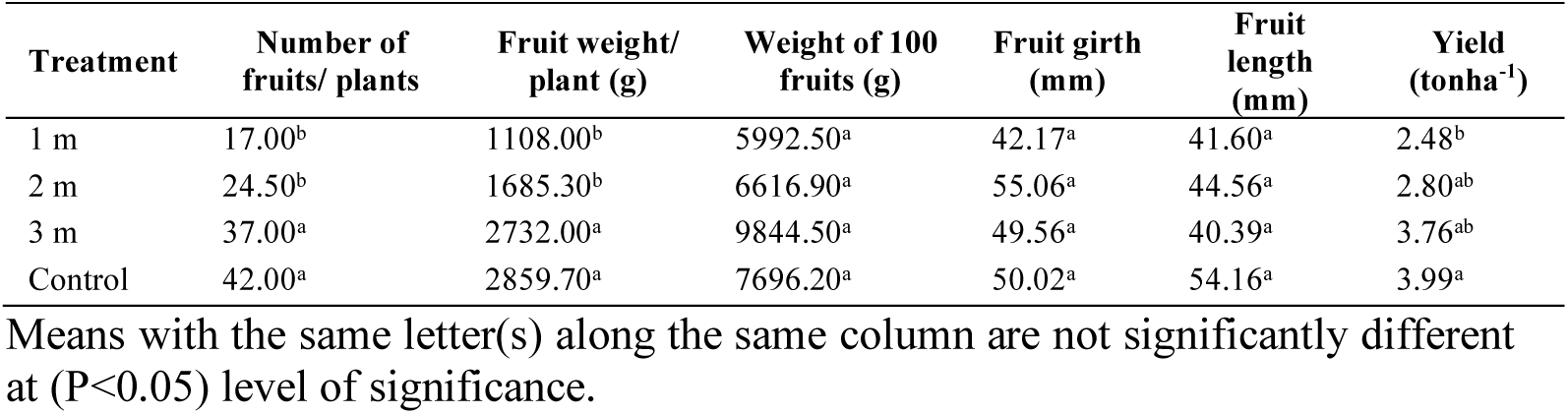
Effect of intercropping on yield components of tomato (NG/AA/SEP/09/053)-juvenile oil palm.

| Treatment | Number of fruits/ plants | Fruit weight/plant (g) | Weight of 100 fruits (g) | Fruit girth (mm) | Fruit length (mm) | Yield (tonha<sup>-1</sup>) |
| --- | --- | --- | --- | --- | --- | --- |
| 1 m | 17.00<sup>b</sup> | 1108.00<sup>b</sup> | 5992.50<sup>a</sup> | 42.17<sup>a</sup> | 41.60<sup>a</sup> | 2.48<sup>b</sup> |
| 2 m | 24.50<sup>b</sup> | 1685.30<sup>b</sup> | 6616.90<sup>a</sup> | 55.06<sup>a</sup> | 44.56<sup>a</sup> | 2.80<sup>ab</sup> |
| 3 m | 37.00<sup>a</sup> | 2732.00<sup>a</sup> | 9844.50<sup>a</sup> | 49.56<sup>a</sup> | 40.39<sup>a</sup> | 3.76<sup>ab</sup> |
| Control | 42.00<sup>a</sup> | 2859.70<sup>a</sup> | 7696.20<sup>a</sup> | 50.02<sup>a</sup> | 54.16<sup>a</sup> | 3.99<sup>a</sup> |

Means with the same letter(s) along the same column are not significantly different at ( $P < 0.05$ ) level of significance.

**TABLE 3:** Interaction of Tomato accessions and intercropping distances on growth parameters and yield components of tomato accessions NGB 01665 and NG/AA/SEP/09/053.

| Factors |  | Number of leaves | Plant height (cm) | Stem girth (mm) | Leaf area (cm<sup>2</sup>) | Chlorophyll index | Total chlorophyll content | Fruit number/plant | Fruit weight / plant (g) | Weight of 100 fruits (g) | Fruit circumference (mm) | Fruit length (mm) | Yield/hectare (tonnes/ha) |
| --- | --- | --- | --- | --- | --- | --- | --- | --- | --- | --- | --- | --- | --- |
| Accessions | Tomato NGB 01665 | 78.594 | 44.832 | 3.793 | 60.97 | 41.662 | 0.026 | 28.25 | 2089 | 8271 | 47.005 | 49.718 | 3.448 |
|  | Tomato NG/AA/SEP/09/053 | 69.875 | 46.991 | 3.252 | 64.378 | 43.852 | 0.029 | 30.125 | 2096 | 7538 | 49.201 | 45.177 | 3.256 |
| Standard error |  | 5.804 | 2.798 | 0.167 | 2.975 | 1.087 | 0.02 | 1.728 | 145.317 | 608.653 | 2.159 | 3.135 | 0.218 |
| Distance | 1 m | 57.312<sup>c</sup> | 35.738<sup>b</sup> | 2.784<sup>b</sup> | 42.806<sup>b</sup> | 41.978<sup>a</sup> | 0.026<sup>a</sup> | 14.625<sup>b</sup> | 99.174<sup>c</sup> | 6212<sup>b</sup> | 41.201<sup>b</sup> | 40.718<sup>b</sup> | 2.578<sup>c</sup> |
|  | 2 m | 62.688<sup>bc</sup> | 45.219<sup>ab</sup> | 3.481<sup>a</sup> | 63.774<sup>a</sup> | 42.967<sup>a</sup> | 0.028<sup>a</sup> | 21.625<sup>b</sup> | 1859<sup>b</sup> | 6898<sup>ab</sup> | 48.825<sup>ab</sup> | 46.731<sup>ab</sup> | 2.911<sup>bc</sup> |
|  | 3 m | 85.062<sup>ab</sup> | 53.875<sup>a</sup> | 3.972<sup>a</sup> | 75.589<sup>a</sup> | 41.238<sup>a</sup> | 0.026<sup>a</sup> | 38.250<sup>a</sup> | 2499<sup>a</sup> | 9346<sup>a</sup> | 50.895<sup>a</sup> | 47.559<sup>ab</sup> | 3.745<sup>ab</sup> |
|  | Control | 91.875<sup>a</sup> | 48.812<sup>a</sup> | 3.854<sup>a</sup> | 68.527<sup>a</sup> | 44.844<sup>a</sup> | 0.031<sup>a</sup> | 42.250<sup>a</sup> | 3016<sup>a</sup> | 9161<sup>a</sup> | 51.491<sup>a</sup> | 54.154<sup>a</sup> | 4.173<sup>a</sup> |
| Standard error |  | 8.208 | 3.957 | 0.237 | 4.208 | 1.537 | 0.02 | 2.444 | 205.509 | 860.765 | 3.053 | 3.135 | 0.309 |
| Interaction | Accession X Distance | * | NS | * | * | NS | NS | NS | NS | NS | NS | NS | NS |

Where “\*” means interaction was significant and “NS” means interaction was not significant for the respective growth and reproductive parameter.

Intercropping is asserted to be one of the most momentous cropping techniques in sustainable agriculture; to its employment a number of environmental advantages, starting with diversification of the outcome of agriculture to promoting land biodiversity. Nchanji *et al*. (2016) opined that this model-intercropping integrates low, medium, and tall plants, as well as plants of short, medium, and long-life cycles, including trees.

**TABLE 4:**
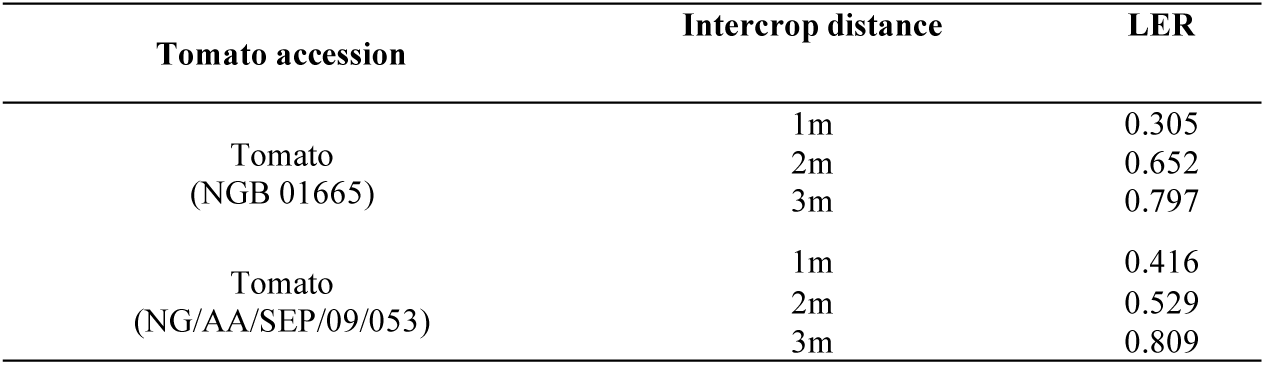
Partial land equivalent ratio (pLER) of tomato-juvenile oil palm intercrops during 2^nd^ year of plantation establishment.

| Tomato accession | Intercrop distance | LER |
| --- | --- | --- |
| Tomato<br>(NGB 01665) | 1m | 0.305 |
|  | 2m | 0.652 |
|  | 3m | 0.797 |
| Tomato<br>(NG/AA/SEP/09/053) | 1m | 0.416 |
|  | 2m | 0.529 |
|  | 3m | 0.809 |

Hutagalung (1995) investigated the intercropping of red pepper, tomato, cabbage and yard-long bean in mango orchard reported that tomato gave high return from intercropping followed by pepper. He concluded that the presence of vegetable crop within young mango trees will contribute additional income to farmers. This indicates that the tomato plants would only tolerate the canopy and perform significantly well if they are intercropped at minimum of 3 m from the juvenile oil palm. Intercropping advantage was not recorded for any of the intercropping distances, with land equivalent ratio values less than unity. This indicates their poor performance when intercropped with juvenile oil palm especially at 1 m and 2 m from juvenile oil palm.

### Effect of intercropping on growth and yield components of tomato (NG/AA/SEP/09/053)

The tomato plants at 3 m from juvenile oil palm recorded a good performance in terms of leaf number, plant height, stem girth and leaf area but the values were not statistically different from those planted in the control plot. Those planted at 1 and 2 m from the juvenile oil palm recorded the poorest performance during the trial. This may be that the tomato could not tolerate the canopy shade from the juvenile oil palm at 1 and 2 m. This observation is in accordance with earlier finding by Olubode *et al.* (2012) who reported growth and yield decline when some fruit vegetables were intercropped with tree crop. They attributed this to competition for light and other developmental resources under the canopy of the tree crop.

Similarly, the tomato plants of the control plot recorded the highest fruit number, fruit weight and yield per hectare during this trial. However, these values recorded were not significantly different from those recorded from tomatoes planted at 3 m from the juvenile oil palm. Sadiq (2011) examined the growth and productivity response of tomato with pepper intercrop and reported high performances of tomatoes in terms of growth and yield when intercropped. The tomato plants intercropped also showed a reduction in yield which could be attributed to the increase in the canopy spread of the juvenile oil palm.

This observation further indicate that tomato plants are sun loving and they will give oil palm plantation farmers significant yield when intercropped at 3 m from the juvenile oil palm. More so, the tomato (NG/AA/SEP/09/053) only recorded LER value close to unity (intercrop advantage) when they were planted at 3 m from juvenile oil palm during the first year.

## CONCLUSION

The responses of the tomato (NGB 01665 and NG/AA/SEP/09/053) to intercropping with juvenile oil palms during the second year of establishment of the oil palm plantation can be successfully practiced. This success in intercrop was achieved by planting the fruit vegetables at a minimum of 2 m from the juvenile oil palm.

However, it was found that intercropping these tomato accessions at 1 m from the juvenile oil palms depressed the performance of the fruit vegetable. This poor performance recorded by the tomato at this distance was probably because light transmission through the juvenile oil palm fronds at this distance in very inadequate to support the growth and yield of the attributes of the tomato accessions.

